# Effects of Bisphenol A on Incidence and Severity of Cardiac Lesions in the NCTR-Sprague-Dawley Rat: A CLARITY-BPA Study

**DOI:** 10.1101/128454

**Authors:** Robin Gear, Jessica A. Kendziorski, Scott M. Belcher

## Abstract

The goal of this study was to determine whether bisphenol A (BPA) had adverse effects indicative of cardiac toxicity. As part of the “*Consortium Linking Academic and Regulatory Insights on BPA Toxicity*” (CLARITY-BPA), study dams and offspring were exposed by daily gavage to five doses of BPA ranging from 2.5 to 25000 μg/kg/day, 0.05 or 0.5 μg/kg/day 17α-ethinyl-estradiol (EE) or 0.3% carboxymethylcellulose vehicle. Exposure-related effects were analyzed in isolated hearts by quantitative morphometry and histopathology. No dose-related changes in body weight were detected. Across all exposure groups including vehicle controls, body weight of continuously dosed males was reduced compared to males dosed only until PND21. Heart weight was increased only in females exposed to EE, and consistent alterations in LV wall thickness were not observed. Exposure-related changes in collagen accumulation were minor and limited to highest EE exposure groups with increased collagen accumulation in PND21 males. Decreased collagen was observed in hearts of BPA or EE exposed females at PND90 and PND180. In BPA or EE treated females cardiomyopathy incidence and severity was significantly increased compared to control females at PND21 with myocardial degeneration observed in both males and females at PND21 and PND90.

## 1. Introduction

Bisphenol A (BPA) is a high volume production chemical used primarily as a monomer in the production of polycarbonate plastics and epoxy resins. The global demand for BPA in 2013 was estimated at more than 7 million metric tons, with demand expected to grow to over 9.6 million metric tons by 2020 (1). Because of its widespread use, and resulting environmental contamination, human exposure to BPA is pervasive. More than a decade ago measurable levels of BPA were detected in 93% of human urine samples analyzed from the 2003-2004 National Health and Nutrition Examination Survey (2). Much concern has been raised about possible adverse health consequences related to these continuous, life-long exposures to BPA.

Results from human association studies investigating health impacts of BPA have been variable and depend on specific study cohort examined and the analysis models used (3). Findings from individual epidemiological studies and a recent systematic review, have however supported an association between higher BPA exposures and increased risk for cardiovascular disease (CVD), obesity, type 2 diabetes, insulin resistance, and hypertension in adults and obesity in children (4-11). Along with human studies associating BPA exposures with CVD, our experimental studies have demonstrated that low nanomolar concentrations of BPA and estrogen (17α-estradiol or E2) could sex-specifically alter estrogen-signaling in cultured adult rodent cardiomyocytes (12). Those effects of BPA were mediated through rapid ERα and ERβ dependent signaling mechanisms that altered Ca^2+^ handling to modify myocyte excitation–contraction coupling. These estrogen-like effects of BPA increased arrhythmia frequencies in response to β-adrenergic stress in isolated hearts from female rats and mice, but not those of males (13). Additional *in vitro* studies have also demonstrated that acute exposures to high concentrations of BPA could decrease the rate and force of contractility and cardiac conduction velocity in hearts from female rats (14, 15) and to a lesser extent in the male heart (16). *In vivo* studies involving analysis of large numbers of male and female CD-1 or C57Bl/6n mice exposed throughout life to a wide range of BPA doses have also identified a number of sex and strain specific exposure-related effects (17-19). Along with having effects on blood pressure, cardiac function, and cardiac adiposity, BPA exposure was also found to alter collagen expression and accumulation in the heart that resulted in abnormal fibrosis and cardiac remodeling. Cardiac transcriptome analysis has also demonstrated that BPA exposures caused sex specific alterations in gene expression that indicated dysregulation of the collagen extracellular matrix and altered lipid metabolism of the heart (17). Those experimental findings supported further the potential for BPA to have negative impacts on heart health, especially in response to cardiac ischemia (17-19).

Although the endocrine disrupting actions of BPA have been exhaustively investigated, there has remained some uncertainty surrounding the potential for BPA to have harmful human health effects. Much of this uncertainty is due to controversies surrounding the design and interpretation of results from hypothesis-driven BPA research studies, and the value of these results for assessing human health risks and regulatory decision making. In an attempt to address these critical uncertainties, an inter-agency collaboration between the National Institute of Environmental Health Sciences’ National Toxicology Program (NIEHS/NTP) and the US Food and Drug Administration (FDA) established the “*Consortium Linking Academic and Regulatory Insights on BPA Toxicity*” (CLARITY-BPA) to perform a comprehensive guideline-compliant 2-year chronic exposure study of the toxicity of BPA (20). The potential impact of the GLP-compliant regulatory toxicity study that formed the backbone for the CLARITY-BPA study was augmented by a parallel study that included a wide-array of investigator initiated hypothesis-driven studies to more comprehensively address the endocrine disrupting actions of BPA. These studies were facilitated by design to leverage the sharing of study tissues to allow analysis of disease-specific endpoints not typically assessed in typical guideline studies of chronic toxicity (20-22). Building these investigator initiated and hypothesis-driven studies around the 2-year chronic toxicity study is anticipated to increase the utility of the investigator-initiated study results for informing hazards characterization and regulatory decision making (20, 21).

In collaboration with all consortium investigators, key aspects of the study design including the doses selected for the CLARITY-BPA study, were agreed upon with the aim of being most useful for all aspects of the study. An over-riding focus was placed on addressing the central regulatory issues of whether chronic BPA exposure results in adverse effects below the current lowest observed effect level. There were five BPA dose groups included in the study with the lowest dose (2.5 μg/kg/day) approaching an estimated human dietary exposure level, and the highest dose (25,000 μg/kg/day) that exceeded the no observed adverse effect level (NOAEL) for systemic toxicity of 5,000 μg/kg/day (23). Along with a vehicle treated control group (aqueous 0.3% carboxymethylcellulose; CMC), there were also two 17α-ethinyl estradiol dose groups (0.05 and 0.5 μg/kg/day) included as comparative controls for effects of an orally bioavailable estrogen. Because many effects of BPA could be developmental, a separate cohort (or “study arm”) of animals dosed only until the time of weaning was also included.

The goal of the presented CLARITY-BPA study analysis was to determine whether BPA had adverse effects on cardiac morphometric and histopathology endpoints indicative of cardiac pathology. The premise for these analyses were derived from findings of our previously published analysis that assessed the structural and functional effects of BPA on the heart in the CD-1 mouse (17). Similar to our previous analysis, exposures in the CLARITY-BPA study occurred through an oral route of administration to mimic a human-relevant route of exposure. Whereas, BPA was incorporated in the ingested rodent diet in the former study, BPA was delivered to the CLARITY-BPA study animals by gavage. Constraints related to complexity of the consortium based study design and animal sharing across all investigator initiated studies did not allow an opportunity for direct assessment of changes in cardiac function or other cardiovascular endpoints of interest (e.g. blood pressure), thus analysis was limited to postmortem morphometric analysis and comparative assessments of cardiac mass and left ventricular (LV) wall thickness, the extent of cardiac fibrosis, and comparative histopathology of the study animal hearts in order to evaluate extents of cardiotoxicity and inflammation induced by each exposure.

## 2. Materials and Methods

### 2.1 Animal Husbandry and Dosing

All animal procedures were performed as a modified guideline-compliant chronic toxicity study that was part of the CLARITY-BPA consortium program (21, 22). Detailed descriptions of study animals, husbandry, randomization procedures, breeding, diet, vehicle, test material preparation and administration, and necropsy are described in detail elsewhere (24). All elements of the experimental design including dose, timing of exposure, and day of sacrifice were developed and agreed upon by the CLARITY-BPA consortium members.

Study animals were maintained on a 12:12 hr light/dark cycle (0600 – 1800) at 23 ± 3 °C with a relative humidity level of 50 ± 20% in an Association for Assessment and Accreditation of Laboratory Animal Care accredited facility. The National Center for Toxicological Research Institutional Animal Care and Use Committee approved all procedures. Approximately 2 weeks prior to breeding, randomly cycling female Sprague-Dawley rats (dams) from the NCTR breeding colony (NCTR-SD strain code 23) were randomized to one of eight exposure groups stratified by body weight to produce approximately equal mean body weights in each group. Dams were mated at 10-14 weeks of age with 11-15 week old males (sires) as previously described with the exception that solid-bottomed polysulfone caging with hardwood chip bedding was used (25). Mating pairs were assigned randomly with the constraint that no sibling or first cousin mating was permitted.

Prior to study assignment dams and sires were fed NIH-41 irradiated pellets (IRR. NIH-41, catalogue #7919C, Harlan Laboratories, Madison, WI) and housed in polycarbonate cages with hardwood chip bedding (P.J. Murphy, Montville, NJ and Lab Animal Supplies, Inc., Lewisville, TX) with water from polycarbonate water bottles. Once assigned to the study at ∼ 21 days of age, breeders (F0) and resulting offspring (F_1_) were housed in polysulfone cages and maintained *ad libitum* on a soy-and alfalfa-free diet (5K96 verified casein diet 10 IF, round pellets, γ-irradiated; Purina Mills, Cat. 1810069) with Millipore-filtered water in glass water bottles with silicone stoppers (#7721 clear, Plasticoid Co., Elkton, MD). Extracts of diet and other study materials were analyzed for BPA, genistein, daidzein, zearalenone, and coumestrol by liquid chromatography and mass spectrometry (25). Each diet lot assayed contained less BPA than the protocol-specified limit of 5 ppb (25), < 1 ppm genistein and daidzein, and < 0.5 ppm zearalenone and coumestrol. Drinking water, polysulfone cage leachates and bedding were also analyzed and found to have BPA levels below the level of the average analytical method blanks (24). After the start of the CLARITY-BPA study, a hypothetical possibility that study animals housed in animal rooms with animals dosed at 250,000 μg/kg/day BPA may have resulted in unintended exposure to low levels of BPA, although there is no direct evidence for contamination of the animals analyzed here (24, 26). Post-analysis sample deidentification revealed no PND90 animals were housed with the 250,000 μg/kg/day BPA animals, 17 of 155 PND21 (Supplemental Table 1), and 240 of 317 PND180 animals (Supplemental Table 2) analyzed were housed in animal rooms with the high BPA exposure group.

Dams and pups were gavaged daily with vehicle (0.3% aqueous CMC, Sigma-Aldrich St. Louis, MO; catalogue C5013, Lot 041M0105V), BPA (CAS 80-05-7, TCI America Portland OR, catalog B0494, Lot 111909/AOHOK, >99% pure) at 2.5 >g BPA/kg bw/day (BPA 2.5), 25.0 >g BPA/kg bw/day (BPA 25), 250 >g BPA/kg bw/day (BPA 250), 2500 >g BPA/kg bw/day (BPA 2500), 25000 >g BPA/kg bw/day (BPA 25000), or 0.05 >g EE/kg bw/day (EE 0.05), and 0.5 >g EE/kg bw/day (EE 0.5). The EE groups were included as an oral bioavailable reference estrogen to establish if specific BPA-related effects were consistent with an “estrogenic” effect. Dose volume was determined immediately after daily body weight collection until 90 days. After 90 days of age dosing was based on weekly body weight. Dosing of dams by gavage was initiated on gestational day (GD) 6 (GD0 = day sperm positive), and continued until day of parturition (PND0). Litters with at least 6 animals were included in the analysis. On PND1 pups were randomly culled from litters with more than 10 animals to achieve a maximum of 5 males and 5 females per litter. Dosing of the F_1_ pups on PND1 by gavage was initiated after litters were culled with daily dosing: 1) continuing until scheduled day of sacrifice at PND21, PND90 (±3 days) or 6 months of age (“continuous dose” groups); or 2) until PND21 with animals housed without dosing until scheduled termination at PND90 (±3 days) or 6 months of age (“stop dose” groups). After weaning the same sex study animals were housed 2 per cage. At PND21, PND 90 and 6 months terminal weights of the F1 animals were collected prior to euthanasia, necropsy and tissue collection.

### 2.2 Tissue Collection, Preparation and Staining

At each of the three time points analyzed (PND21 (weaning), PND90, or 6 months), animals were weighed, sacrificed and hearts were harvested with heart weights recorded at NCTR. Each tissue specimen and all corresponding experimental endpoints (e.g. body and heart weight measurements) were assigned a coded identification and all tissue preparation and subsequent histopathologic analysis and scoring was done blinded to exposure group, exposure duration or sex. Hearts were fixed for 24 hours in 10% formalin, post-fixed in fresh neutral buffered formalin for an additional 24 hours, and then transferred to 70% ethanol for shipping. Hearts were shipped to the Belcher laboratory with these coded identification numbers to ensure that the investigators were completely blinded to all experimental variables except age. Upon arrival, specimens were washed in 70% ethanol, prepared by automated tissue processing for 40-45 minutes each in 7 changes of graded alcohols followed by embedding with 3 changes in paraffin at 58°C with applied vacuum (Tissue-Tek VIP 3000; Sakura Torrence, CA). Hearts were then cut into concentric 1mm transverse sections, and embedded into paraffin blocks (Histocenter 3; Thermo-Shandon Kalamazoo, MI). Serial 5 >m microtome sections were cut from these blocks at 4°C, placed on positively charged slides for staining and analyzed as described previously (17, 27). Heart sections were stained with hematoxylin and eosin (H&E; Richard-Allan, Kalamazoo, MI) using a standard protocol to examine tissue structure, morphology and pathology. Left ventricular free wall area was measured and the average LV free wall thickness was calculated from a single section at the level of the papillary muscle for each study animal. Serial transverse sections at the level of the papillary muscle were also stained with Picrosirius Red (Polysciences; Warrington, PA) to visualize total collagen (red) with bright field illumination (28-31). For picrosirius red staining, tissue sections were deparaffinized, rehydrated, and stained for 8 minutes with Weigert’s hematoxylin (American MasterTech; Lodi, CA). Stained sections were rinsed with tap water for 5 minutes, incubated for 2 minutes in 0.2% phosphomolybdic acid hydrate, and rinsed in deionized H_2_O for 30 seconds. Slides were then stained for 1 hour in picrosirius red F3B solution (1.3% 2,4,6-trinitrophenol, 0.4% Direct Red 80), transferred to 0.1 N hydrochloric acid solution for 2 minutes, washed in 70% ethanol for 45 seconds, dehydrated and then coverslipped.

Stained sections were examined on Nikon Eclipse 55i and 80i microscopes equipped with DS-Fi1 CCD cameras controlled by Digital Sight Software (Nikon; Melville, NY). Digital images of each section were collected using a 1x and 10x objectives, with additional higher magnification images collected using 20x and 40x objectives. Acquired images of picrosirius red sections were captured in RGB file format and then converted to HSI file format using Image Pro v4.5 (Media Cybernetics Silver Springs, MD). Images were confirmed as not containing saturated pixels, and threshholded to background staining intensity. Total left ventricle (LV) area and the total LV collagen staining were calculated with collagen staining reported as a percent of LV area. Visual inspection of each slide was performed to qualitatively confirm the accuracy of the computed levels of staining. From the H&E stained sections, 10x digital images were acquired to examine gross and microscopic tissue structure and to measure LV free wall thickness. The average free LV wall thickness, LV diameter, and total LV area were measured as described previously with Image-Pro v4.5 (17, 27). For wall thickness measurements, 5 evenly spaced digital lines spanning the width of the left ventricular free wall were superimposed onto the digital image from a single stained section at the level of the papillary muscle. The average LV wall thickness was calculated as the mean length of those five lines relative to a stage micrometer of known length. For morphometric data collection and analysis, all samples including controls were comprehensively masked and were analyzed by a single observer blind to exposure dose, exposure duration (stop vs. continuous) and sex (32). All digital results were confirmed by direct microscopic observation.

### 2.3 Evaluation of Cardiac Pathology

Pathology of PND21, PND90, and 6 month deidentified specimens was evaluated by examination of H&E stained sections at final magnifications of 100-200x. Cardiomyopathy, late stage cardiomyopathy (focal fibrosis), diffuse degeneration, and inflammatory infiltration phenotypes were each scored according to the Standardized System of Nomenclature and Diagnostic Criteria (SSNDC) Guide (33). No threshold for morphological changes was applied to the analysis and any lesion consistent with each pathology was scored as positive. Cardiomyopathy and LV pathology was assessed using a standardized four point severity scale employed by Jokinen et al (34, 35) with 1 = <10%, 2 = 11-40%, 3 = 41-80%, 4 = >81% of LV area involvement. Specimens containing visible hemosiderin were also noted. In consultation with a board-certified veterinary pathologist (Diplomat of the American College of Veterinary Pathologists), all pathology was assessed by the same investigator (RG). An independent blinded pathology review was performed by a second investigator (JK) with any differences in lesion grading reviewed and resolved by consensus of the research team.

### 2.4 Data Coding and Decoding Procedures

Details of the data tracking, coding, decoding and quality assurance procedures are described elsewhere (24). To avoid potential bias all primary data collection and analysis was conducted blind to exposure and sex. All individual samples were received with a unique numeric identifier assigned by the NCTR staff. Upon completion of primary data collection, analysis and data quality review (SB and RG), the coded primary data were submitted to the NTP Chemical Effects in Biological Systems (CEBS) data base administrator. Data was then independently verified to contain all expected data for each endpoint, and upon approval of the data Decoding Team all files were “locked” such that data could not be altered (read only format). Upon archiving of data from all University-based research projects which shared a common code, the data decoding information was supplied by NCTR to the CEBS Administrator who performed quality assurance reviews of data integrity and the decoding information. Upon approval by the CLARITY-BPA Decoding Team, the verified decoding information was supplied to the PI (SMB).

### 2.5 Statistical Analysis

Detailed review and analysis of all the research plans included in the CLARITY-BPA hypothesis driven studies followed NIEHS and NTP recommendation for evaluation of 10 animals per sex per group. Our previous analysis had found that n = 10 was a sufficient sample size with enough statistical power to detect BPA exposure induced changes in the CD-1 mouse heart for endpoints assessed here (17). Confound from litter effects are avoided by limiting analysis at each time point to one animal of a given sex from each litter, with each sex being considered separately. The statistical unit used was the litter for all analyses. A minimal level of statistical significance for differences in values among or between groups was considered p < .05. All statistical analyses for differences in values compared to control were made independently for BPA and EE exposures and followed guidelines for low dose endocrine disrupting chemicals (36). Percentage data was arcsine transformed (arcsine of the square root of the value) prior to statistical analysis. Data analysis was performed using Dunnett’s multiple comparison tests, one-way or two-way analysis of variance as indicated, and for pathology severity scores, a rank order ANOVA Kruskal-Wallis H test with Dunn's multiple comparisons tests were used. All data was analyzed using Excel (Microsoft; Redmond, WA) and GraphPad Prism^®^ v6 software (GraphPad; La Jolla, CA).

## 3. Results

### 3.1 Morphometric Characterization: Body Weight

Analysis of variance showed there were no exposure related changes in body weight detected at any time point analyzed (Table 1). An effect of exposure duration (stop dose vs. continuous dose) on male body weights at PND90 and 6 months was observed. Across all exposure groups a two factor analysis of variance (exposure, dose duration) indicated a significant effect of dose duration at PND 90 [*F*(1, 135) = 5.3, *p=* .02], and at 6 months [*F*(1, 135) = 25.0, *p* < .0001]. At PND90 mean body weight of stop dose control males was 9.5% greater than in the continuously exposed control group (stop dose *M* = 490, SD = 43.1; continuous dose *M* = 464, SD = 38.3). Similarly at 6 month mean body weight of stop dose control males was 9.1% greater than in the continuously exposed control group (stop dose *M* = 644, SD = 69.5; continuous dose *M* = 583, SD = 40.5). Female body weight was not influenced by exposure duration at either time point.

**Table 1.**
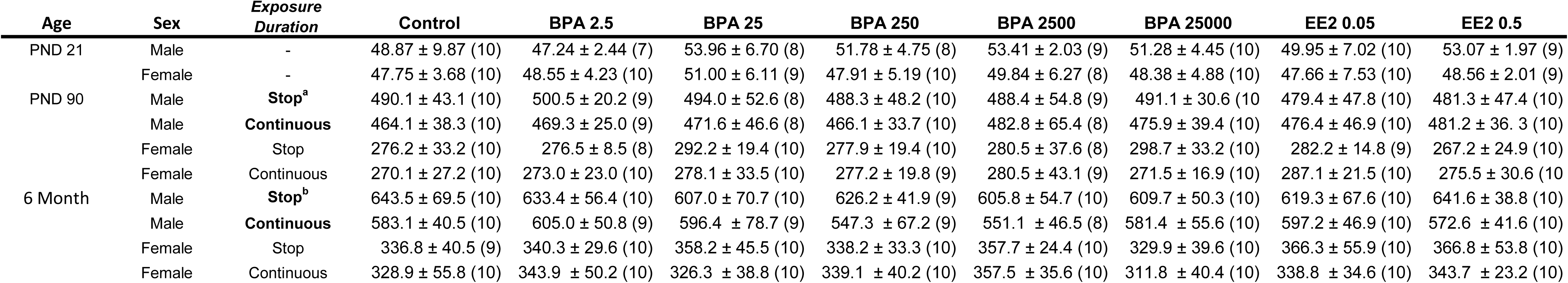
Male and Female Body Weight (g)

### 3.2 Morphometric Characterization: Heart Weight

At PND21 analysis of variance showed there was no main effect on absolute heart weight [BPA male: *F*(5, 48) = 0.60, *p* = .70; female *F*(5, 52) = 1.08, *p* = .38: EE male *F*(2, 27) = 1.84, *p* = .10; female *F*(2, 26) = 0.68, *p* = 0.52] or relative heart weight indexed to body weight [BPA male: *F*(5, 47) = 0.17, *p* = .97; female *F*(5, 51) = 0.87, *p* = .51: EE male *F*(2, 26) = 0.56, *p* = .58; female *F*(2, 26) = 0.76, *p* = .48]. At PND90 absolute heart weight was influenced by exposure in females continuously exposed to EE (Table 2). Dunnett’s multiple comparison tests found at PND90 chronic the heart weight in females from the continuous 0.05 EE (*p* = .004, *d* = 2.03) and 0.5 EE (*p =* .0003, *d* = 2.12) exposure groups were significantly increased compared to control. When heart weight at PND90 was indexed to body weight (Table 3), a significant effect was identified in both the 0.5 EE stop dose (*p* = .013, *d* = 1.48) and 0.5 EE continuous dose groups (*p* = .012, *d* = 1.27). At 6 months of age heart weight was significantly decreased in stop dose 2.5 BPA females (*p* = .02, *d* = 1.38), and although the decrease in indexed heart weight for the stop dose 2.5 BPA females failed to reach significance (*p* = .06), the effect size remained large (*d* = 1.11). Heart weight (*p* = .001, *d* = 1.61) and indexed heart weight (*p* = .0009, *d* = 1.88) were significantly increased in the 0.5 EE continuously dosed females. An increase in the mean indexed heart weight for females in the continuously dosed 25 BPA group was noted, but did not reach the criteria set for statistical significance, although the effect size was again large (*p* = .053; *d* = 0.88). No other changes in absolute (Table 2) or indexed heart weight (Table 3) were identified.

**Table 2.**
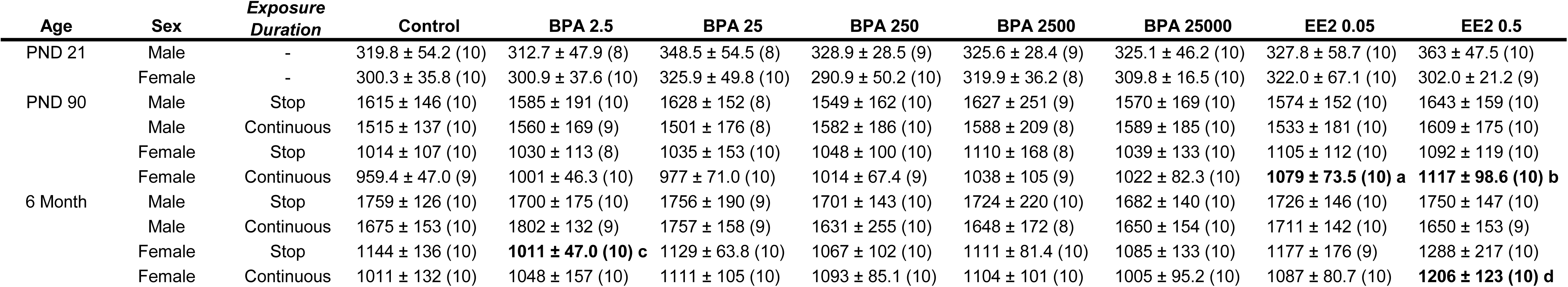
Male and Female Heart Weight (mg)

**Table 3.** Male and Female Heart Weight/Body Weight (mg/g)

| Age | Sex | Exposure Duration | Control | BPA 2.5 | BPA 25 | BPA 250 | BPA 2500 | BPA 25000 | EE2 0.05 | EE2 0.5 |
| --- | --- | --- | --- | --- | --- | --- | --- | --- | --- | --- |
| PND 21 | Male | - | 6.38 ± 0.65 (9) | 6.35 ± 0.51 (8) | 6.46 ± 0.71 (8) | 6.37 ± 0.33 (8) | 6.20 ± 0.65 (10) | 6.34 ± 0.66 (10) | 6.55 ± 0.68 (10) | 6.67 ± 0.66 (10) |
|  | Female | - | 6.16 ± 0.27 (9) | 6.19 ± 0.40 (10) | 6.35 ± 0.73 (10) | 6.04 ± 0.57 (10) | 6.44 ± 0.54 (8) | 6.45 ± 0.66 (10) | 6.43 ± 0.69 (9) | 6.23 ± .041 (10) |
| PND 90 | Male | Stop | 3.30 ± 0.19 (10) | 3.19 ± 0.37 (10) | 3.31 ± 0.27 (8) | 3.17 ± 0.15 (10) | 3.33 ± 0.38 (9) | 3.20 ± 0.27 (10) | 3.29 ± 0.19 (10) | 3.43 ± 0.34 (10) |
|  | Male | Continuous | 3.27 ± 0.23 (10) | 3.32 ± 0.19 (9) | 3.18 ± 0.26 (8) | 3.40 ± 0.33 (10) | 3.30 ± 0.30 (8) | 3.34 ± 0.23 (10) | 3.22 ± 0.27 (10) | 3.34 ± 0.21 (10) |
|  | Female | Stop | 3.69 ± 0.29 (10) | 3.72 ± 0.37 (8) | 3.55 ± 0.17 (10) | 3.78 ± 0.36 (10) | 3.96 ± 0.22 (8) | 3.48 ± 0.13 (10) | 3.92 ± 0.34 (10) | **4.09 ± 0.29 (10) a** |
|  | Female | Continuous | 3.67 ± 0.37 (10) | 3.68 ± 0.18 (10) | 3.54 ± 0.32 (10) | 3.46 ± 0.38 (10) | 3.76 ± 0.57 (9) | 3.76 ± 0.21 (10) | 3.77 ± 0.24 (10) | **4.07 ± 0.29 (10) b** |
| 6 Month | Male | Stop | 2.75 ± 0.19 (10) | 2.69 ± 0.19 (10) | 2.90 ± 0.21 (9) | 2.79 ± 0.24 (10) | 2.84 ± 0.18 (10) | 2.77 ± 0.18 (10) | 2.80 ± 0.21 (10) | 2.73 ± 0.25 (10) |
|  | Male | Continuous | 2.87 ± 0.20 (10) | 2.99 ± 0.27 (9) | 2.97 ± 0.25 (9) | 2.85 ± 0.44 (10) | 2.99 ± 0.11 (8) | 2.85 ± 0.26 (10) | 2.87 ± 0.11 (10) | 2.89 ± 0.23 (9) |
|  | Female | Stop | 3.28 ± 0.33 (10) | **2.99 ± 0.21 (10) c** | 3.18 ± 0.31 (10) | 3.16 ± 0.22 (10) | 3.10 ± 0.12 (10) | 3.30 ± 0.30 (10) | 3.20 ± 0.31 (9) | 3.52 ± 0.39 (10) |
|  | Female | Continuous | 3.09 ± 0.26 (10) | 3.06 ± 0.26 (10) | **3.45 ± 0.54 (10) e** | 3.24 ± 0.28 (10) | 3.10 ± 0.19 (10) | 3.24 ± 0.19 (10) | 3.22 ± 0.23 (10) | **3.50 ± 0.19 (10) d** |

### 3.3 LV Wall Thickness and Fibrosis

At PND 21 an analysis of variance showed there were no main effects of either BPA or EE on LV wall thickness in either sex (Table 4). At PND90 Dunnett’s multiple comparison tests indicated a significant decrease in LV wall thickness in the female stop dose 2.5 BPA group only (*p* = .049, *d* = 1.19). In the 0.05 EE continuous dose group at 6 months a significant (*p* = .048, *d* = 1.03) increase in LV wall thickness of females was identified (Table 4).

**Table 4.** Male and Female LV Wall thickness (mm)

| Age | Sex | Exposure<br>Duration | Control | BPA 2.5 | BPA 25 | BPA 250 | BPA 2500 | BPA 25000 | EE2 0.05 | EE2 0.5 |
| --- | --- | --- | --- | --- | --- | --- | --- | --- | --- | --- |
| PND 21 | Male | - | 1.85 ± 0.17 (10) | 1.81 ± 0.15 (8) | 1.74 ± 0.11 (8) | 1.90 ± 0.32 (9) | 1.89 ± 0.14 (10) | 1.99 ± 0.32 (10) | 1.88 ± 0.22 (10) | 1.95 ± 0.32 (10) |
|  | Female | - | 1.83 ± 0.18 (10) | 1.91 ± 0.21 (10) | 1.85 ± 0.10 (10) | 1.84 ± 0.16 (10) | 1.89 ± 0.12 (10) | 1.87 ± 0.20 (10) | 1.74 ± 0.25 (10) | 1.72 ± 0.16 (10) |
| PND 90 | Male | Stop | 2.57 ± 0.44 (10) | 2.64 ± 0.46 (10) | 2.46 ± 0.22 (8) | 2.49 ± 0.23 (10) | 2.44 ± 0.31 (9) | 2.48 ± 0.28 (10) | 2.57 ± 0.37 (10) | 2.37 ± 0.27 (10) |
|  | Male | Continuous | 2.58 ± 0.23 (10) | 2.37 ± 0.28 (9) | 2.38 ± 0.28 (8) | 2.38 ± 0.20 (10) | 2.52 ± 0.45 (8) | 2.45 ± 0.22 (10) | 2.49 ± 0.27 (10) | 2.43 ± 0.26 (10) |
|  | Female | Stop | 2.60 ± 0.29 (10) | **2.30 ± 0.22 (8) a** | 2.49 ± 0.20 (10) | 2.47 ± 0.28 (10) | 2.49 ± 0.20 (8) | 2.49 ± 0.22 (10) | 2.55 ± 0.23 (10) | 2.59 ± 0.30 (10) |
|  | Female | Continuous | 2.36 ± 0.22 (10) | 2.53 ± 0.39 (10) | 2.28 ± 0.26 (10) | 2.40 ± 0.17 (10) | 2.46 ± 0.30 (9) | 2.22 ± 0.22 (10) | 2.47 ± 0.26 (10) | 2.51 ± 0.18 (10) |
| 6 Month | Male | Stop | 2.81 ± 0.43 (10) | 2.68 ± 0.20 (10) | 2.66 ± 0.38 (9) | 2.72 ± 0.28 (10) | 2.53 ± 0.29 (10) | 2.75 ± 0.25 (9) | 2.46 ± 0.18 (10) | 2.43 ± 0.22 (10) |
|  | Male | Continuous | 2.54 ± 0.30 (10) | 2.73 ± 0.43 (9) | 2.62 ± 0.37 (9) | **2.95 ± 0.47 (10)c** | 2.69 ± 0.38 (8) | 2.65 ± 0.32 (10) | 2.61 ± 0.25 (10) | 2.60 ± 0.36 (9) |
|  | Female | Stop | 2.44 ± 0.19 (10) | 2.34 ± 0.18 (10) | 2.40 ± 0.25 (10) | 2.68 ± 0.27 (10) | 2.73 ± 0.41 (10) | 2.49 ± 0.30 (10) | 2.44 ± 0.33 (10) | 2.62 ± 0.24 (10) |
|  | Female | Continuous | 2.28 ± 0.36 (10) | 2.39 ± 0.27 (10) | 2.45 ± 0.33 (10) | 2.38 ± 0.26 (10) | 2.51 ± 0.24 (10) | 2.49 ± 0.23 (10) | **2.63 ± 0.35 (10)b** | 2.44 ± 0.27 (9) |

Collagen accumulation was similar to control in each exposure group for both males and females at PND21 (Table 5) with Dunnett’s multiple comparison tests revealing a significant increase (*p* = .027; *d* = 1.15) of LV collagen in male hearts from the 0.5 EE exposure group (Table 5). For males at PND90 a two factor analysis of variance (exposure, duration) indicated a significant effect of both exposure [*F*(1, 133) = 2.27, *p* = .033] and dose duration [*F*(1, 133) = 14.5, *p* = .0002] on amount of LV collagen that was not qualified by an interaction [*F*(7, 133) = 1.59, *p* = .14]. The mean percentage of LV collagen in continuously dosed control males (*M* = 3.38, SD = 1.18) was significantly increased (*t*(18) = 2.37, *p* = .029, *d* = 1.12) compared to stop dose male control (*M* = 2.34, SD = 0.86). In females at PND90 LV collagen was not influenced by exposure or dose duration. There was no discernable effect of dose duration on collagen accumulation in either sex at the 6 month time point, and Dunnett’s multiple comparison tests identified a significant (*p* = .006, *d* = 1.37) decrease of collagen in female hearts from the stop dose 0.5 EE group (Table 5).

**Table 5.** Male and Female Fibrosis (percentage of LV Area)

| Age | Sex | Exposure Duration | Control | BPA 2.5 | BPA 25 | BPA 250 | BPA 2500 | BPA 25000 | EE2 0.05 | EE2 0.5 |
| --- | --- | --- | --- | --- | --- | --- | --- | --- | --- | --- |
| PND 21 | Male | - | 0.94 ± 0.33 (10) | 1.37 ± 0.37 (8) | 1.33 ± 0.66 (8) | 0.98 ± 0.42 (9) | 1.12 ± 0.43 (10) | 0.99 ± 0.26 (10) | 1.13 ± 0.33 (10) | **1.40 ± 0.55 (10)a** |
|  | Female | - | 1.14 ± 0.55 (10) | 1.09 ± 0.27 (10) | 1.16 ± 0.43 (10) | 1.11 ± 0.45 (10) | 1.32 ± 0.42 (10) | 1.16 ± 0.30 (10) | 1.22 ± 0.49 (10) | 1.02 ± 0.38 (10) |
| PND 90 | Male | Stop | 2.34 ± 0.86 (10) | 2.32 ± 0.54 (9) | 2.63 ± 0.81 (8) | 3.04 ± 1.08 (10) | 3.23 ± 1.05 (9) | 2.90 ± 1.06 (10) | 2.61 ± 0.60 (10) | 2.81 ± 0.61 (9) |
|  | Male | Continuous | 3.38 ± 1.18 (10) | 3.09 ± 0.96 (9) | 3.44 ± 0.93 (8) | 3.09 ± 1.19 (10) | 4.18 ± 1.06 (8) | 2.36 ± 0.39 (9) | 3.41 ± 1.08 (10) | 3.65 ± 1.10 (10) |
|  | Female | Stop | 2.70 ± 1.34 (10) | 2.46 ± 0.88 (8) | 2.64 ± 1.1 (10) | 2.59 ± 0.72 (10) | 3.16 ± 1.32 (8) | 3.11 ± 1.02 (10) | 2.01 ± 0.41 (10) | 2.31 ± 0.57 (10) |
|  | Female | Continuous | 3.13 ± 0.90 (10) | 2.74 ± 1.21 (10) | 2.81 ± 0.82 (9) | 3.13 ± 1.02 (10) | 3.16 ± 0.97 (9) | **2.14 ± 0.53 (9)b** | 2.88 ± 1.01 (10) | 3.00 ± 1.13 (10) |
| 6 Month | Male | Stop | 2.50 ± 0.82 (10) | 2.19 ± 0.89 (10) | 1.88 ± 0.52 (9) | 2.19 ± 0.88 (10) | 2.47 ± 0.67 (9) | 2.35 ± 0.75 (10) | 2.60 ± 0.64 (10) | 2.54 ± 0.99 (10) |
|  | Male | Continuous | 2.48 ± 0.77 (10) | 2.29 ± 0.59 (9) | 2.50 ± 1.00 (10) | 2.45 0.93 (10) | 2.69 ± 0.79 (8) | 3.28 ± 1.40 (10) | 2.18 ± 0.43 (9) | 2.30 ± 0.42 (8) |
|  | Female | Stop | 2.82 ± 0.83 (10) | 2.99 ± 1.10 (10) | 2.88 ± 1.08 (10) | 2.58 ± 1.14 (10) | 2.31 ± 0.68 (10) | 2.96 ± 0.68 (9) | 2.23 ± 0.43 (9) | **1.90 ± 0.60 (10)c** |
|  | Female | Continuous | 2.31 ± 1.08 (10) | 2.47 ± 0.96 (10) | 2.35 ± 0.89 (10) | 2.17 ± 0.43 (10) | 2.83 ± 1.01 (10) | 2.69 ± 0.90 (10) | 2.59 ± 0.92 (10) | 2.47 ± 1.02 (10) |

### 3.4 Histopathology: Progressive Cardiomyopathy (PCM)

Cardiomyopathy-like lesions were frequently observed in the sections used for characterization of LV wall thickness and fibrosis. As a result the incidence and severity of myocardial lesions were characterized to investigate the hypothesis that there was a high background level of cardiomyopathy in the hearts of control, and that exposure to EE or BPA was increasing the incidence and severity of these lesions. Analysis of single transverse sections of the heart for each animal identified lesions in 90% of males and 60% of control females at PND21 (Table 6). In BPA or EE treated females at PND21 cardiomyopathy incidence was increased compared to control females and a significant increase in severity was found for 2.5, 250, 25,000 μg/kg/day BPA and each EE group (Table 6). In a male exposed to 250 μg/kg/day BPA and a female from each of the two lowest BPA dose groups (2.5 and 25 μg/kg/day) a diffuse degeneration phenotype involving much of the myocardium was also observed. Shown in 1 are photomicrographs of H&E stained hearts sections from control females showing representative lesions observed at PND21. Small regions of inflammatory cell infiltrates indicative of the earliest stages of cardiomyopathy were frequently noted (Fig. 1A arrows). Areas of myocyte degeneration with extensive vacuolation of myocytes and lacking evident fibrosis were also observed (Fig. 1B). Regions of highly disorganized myocyte morphology, with evident myocyte degeneration, diffuse vacuolation, increased cellularity and fibrosis consistent with a diagnosis of mid-stage PCM were readily apparent (Fig. 1C). At this age regions of focal fibrosis (late stage cardiomyopathy), often associated with the endocardium, were also identified in the LV myocardium and the papillary muscle (Fig. 1D).

**Figure 1.**
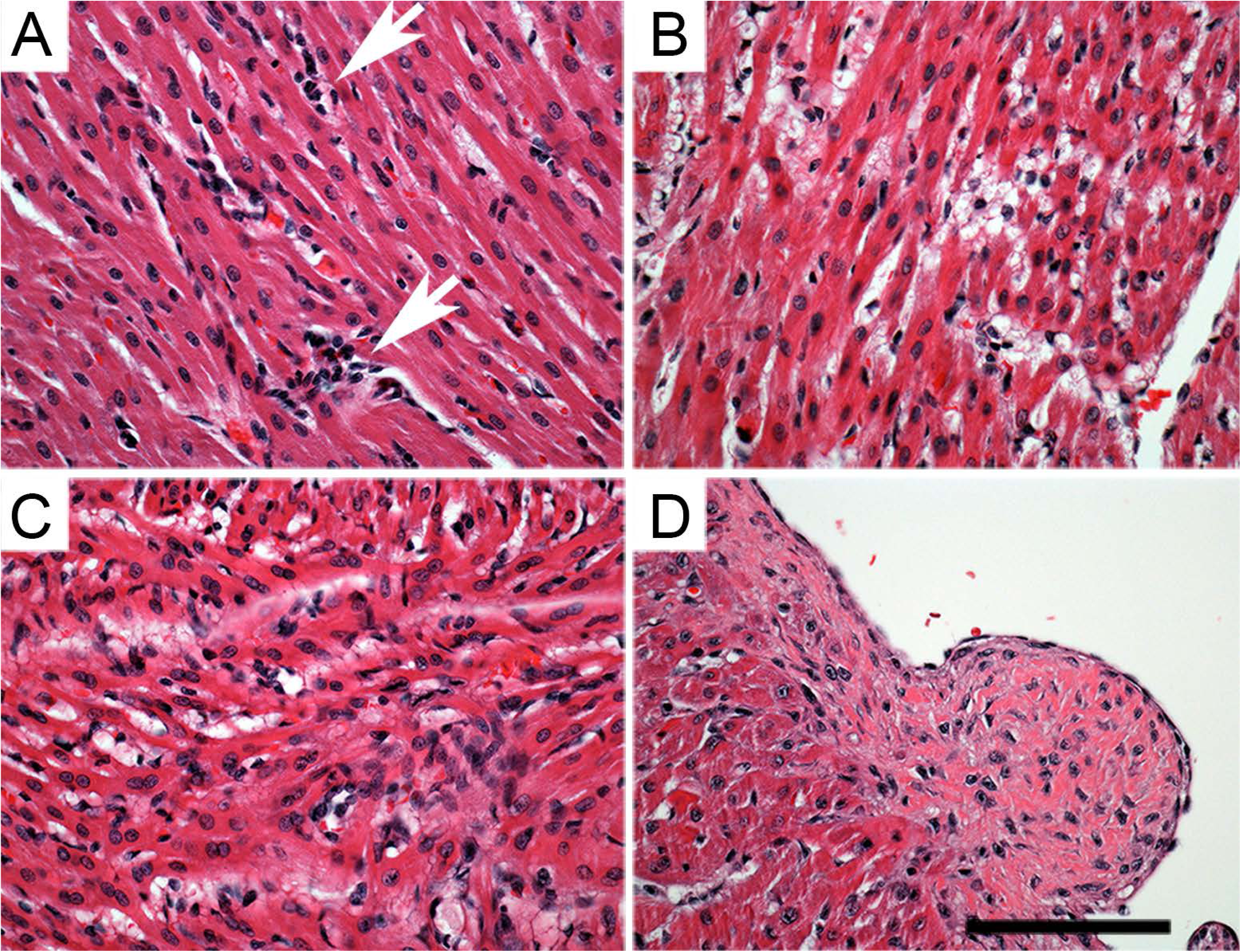
**Cardiomyopathy at PND21.** Small regions of inflammatory cell infiltrates (white arrows) indicative of the earliest stages of cardiomyopathy (A). Areas of myocyte degeneration with extensive vacuolation of myocytes and lacking evident fibrosis (B). Small regions of mid-stage PCM with highly disorganized myocyte morphology, evident myocyte degeneration, diffuse vacuolation, increased cellularity and fibrosis (C). Focal fibrosis indicative of late stage cardiomyopathy (D). Bars = 100 μm

**Table 6.**
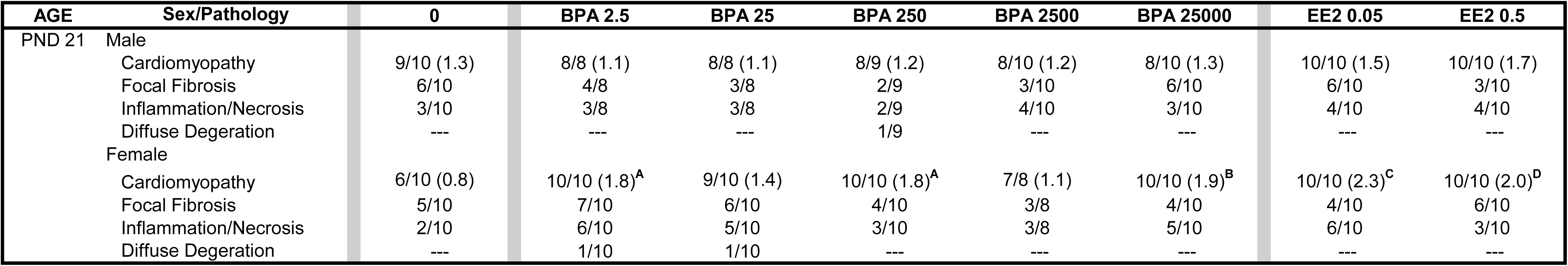
Incidence of Cardiac Lesions at PND 21

At PND90 (Table 7) and 6 months (Table 8) cardiomyopathy in both males and females was observed in 100% of control samples from both the stop dose and the continuous dose arms of the study. Cardiomyopathy incidence at PND90 was essentially quantitative across all dose groups with lesion severity similar for each exposure group. A diffuse degeneration phenotype at PND90 (Table 7) was observed in males and females from the continuous (males: BPA 25, 250, 25,000; EE 0.5 μg/kg/day; females: BPA 2.5, 25; EE 0.05, 0.5 μg/kg/day) and stop dose (males: BPA 250, 25,000; EE 0.05, 0.5 μg/kg/day; Females 250 μg/kg/day) exposure groups. Cardiomyopathy at this age was characterized by degenerating myofibrils and associated focal and multifocal inflammatory cell infiltrates (Figure 2A). In some cases hemosiderin containing macrophages suggestive of previous vascular hemorrhage were observed (Fig. 2B; arrowheads). Regions of focal fibrosis indicative of late-stage cardiomyopathy were characteristic of both control females (Fig. 2C) and males (Fig. 2D) and each of the exposure group. At both PND90 (Tables 7) and 6 months (Table 8) the incidence of the inflammatory phenotype was greater in control males than in females, and lesions were more often multifocal with a larger area of involvement (Figure 2E). In some cases myocyte degeneration, necrosis, and inflammation with hemosiderin containing macrophages were observed (Fig. 2F; arrows). Exposure related effects on cardiomyopathy above the background observed in controls were not detectable at this later time point.

**Figure 2.**
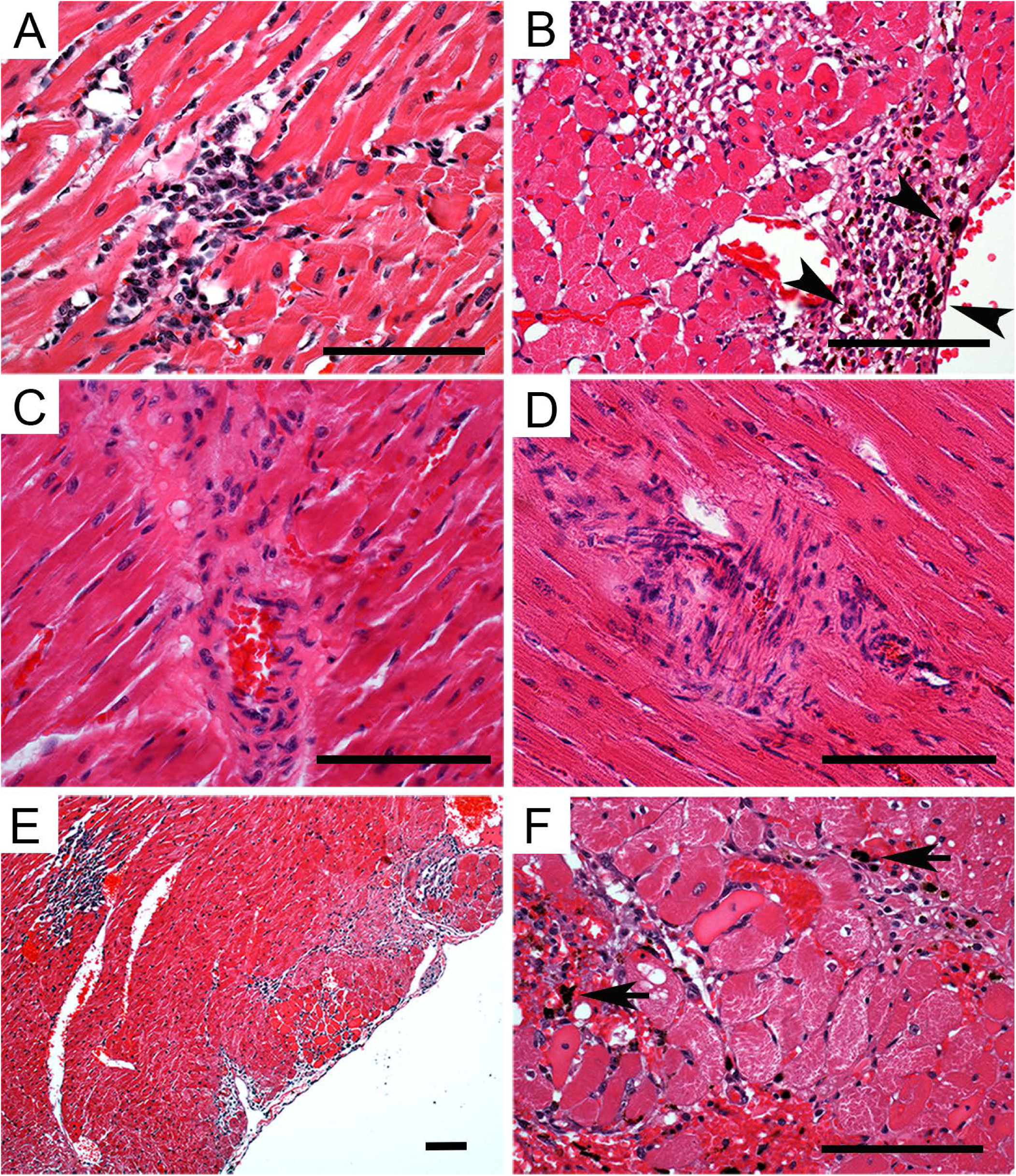
**Cardiomyopathy at PND90 and 6 months**. Lesions at PND90 were characterized by degenerating myofibrils associated focal and multifocal inflammatory cell infiltrates (A). Regions of extensive myocyte necrosis with inflammatory cell infiltrates and hemosiderin containing macrophages (B; arrowheads). Focal perivascular and interstitial fibrosis indicative of late-stage cardiomyopathy in control female (C) and male (D) at PND90. At 6 months cardiomyopathy at this age was characterized by multifocal lesions with a larger area of involvement (E). Evident degenerating myofibrils associated focal and multifocal inflammatory cell infiltrates with hemosiderin containing macrophages (arrows) suggestive of previous vascular hemorrhage were also observed (F). Bars = 100 μm.

**Table 7.**
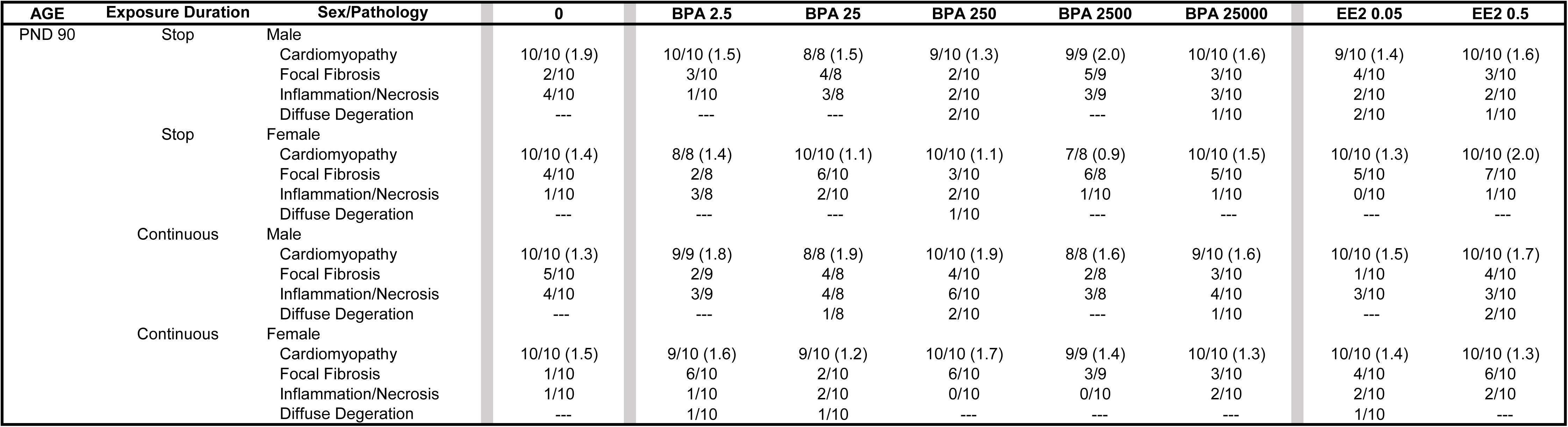
Incidence of Cardiac Lesions at PND90

**Table 8.** Incidence of Cardiac Lesions at 6 Months

| AGE | Exposure Duration | Sex/Pathology | 0 | BPA 2.5 | BPA 25 | BPA 250 | BPA 2500 | BPA 25000 | EE2 0.05 | EE2 0.5 |
| --- | --- | --- | --- | --- | --- | --- | --- | --- | --- | --- |
| 6 Month | Stop | Male (potential exposure)* | 10/10 | 10/10 | 8/9 | 4/10 | 10/10 | 10/10 | 6/10 | 8/10 |
|  |  | Cardiomyopathy | 10/10 (1.8) | 10/10 (1.9) | 9/9 (1.9) | 10/10 (1.7) | 10/10 (1.9) | 10/10 (1.6) | 10/10 (1.5) | 10/10 (1.8) |
|  |  | Focal Fibrosis | 6/10 | 4/10 | 4/9 | 5/10 | 6/10 | 5/10 | 5/10 | 4/10 |
|  |  | Inflammation/Necrosis | 6/10 | 9/10 | 6/9 | 5/10 | 6/10 | 7/10 | 5/10 | 6/10 |
|  |  | Diffuse Degeration | --- | --- | --- | --- | --- | --- | --- | --- |
|  | Stop | Female (potential exposure)* | 10/10 | 10/10 | 8/10 | 10/10 | 8/10 | 8/10 | 8/10 | 10/10 |
|  |  | Cardiomyopathy | 10/10 (1.6) | 10/10 (1.6) | 9/10 (1.7) | 8/10 (1.2) | 10/10 (1.6) | 9/10 (1.4) | 9/10 (1.8) | 10/10 (1.9) |
|  |  | Focal Fibrosis | 3/10 | 4/10 | 3/10 | 3/10 | 7/10 | 1/10 | 5/10 | 9/10 |
|  |  | Inflammation/Necrosis | 0/10 | 1/10 | 4/10 | 3/10 | 1/10 | 2/10 | 3/10 | 4/10 |
|  |  | Diffuse Degeration | --- | --- | --- | --- | --- | --- | --- | --- |
|  | Continuous | Male (potential exposure)* | 3/10 | 2/10 | 0/10 | 2/10 | 2/8 | 2/10 | 6/10 | 8/10 |
|  |  | Cardiomyopathy | 10/10 (2.0) | 9/9 (2.1) | 8/9 (1.6) | 10/10 (2.0) | 8/8 (2.1) | 10/10 (1.7) | 10/10 (1.7) | 9/9 (1.8) |
|  |  | Focal Fibrosis | 4/10 | 3/9 | 2/9 | 6/10 | 3/8 | 2/10 | 3/10 | 0/9 |
|  |  | Inflammation/Necrosis | 9/10 | 7/9 | 4/10 | 4/8 | 4/8 | 8/10 | 5/10 | 5/9 |
|  |  | Diffuse Degeration | --- | --- | --- | --- | --- | --- | --- | --- |
|  | Continuous | Female (potential exposure)* | 10/10 | 10/10 | 8/10 | 10/10 | 8/10 | 8/10 | 6/10 | 10/10 |
|  |  | Cardiomyopathy | 10/10 (1.8) | 8/10 (1.4) | 10/10 (1.6) | 10/10 (1.9) | 10/10 (1.8) | 10/10 (2.3) | 10/10 (1.9) | 8/10 (1.6) |
|  |  | Focal Fibrosis | 5/10 | 3/10 | 2/10 | 6/10 | 5/10 | 5/10 | 4/10 | 4/10 |
|  |  | Inflammation/Necrosis | 0/10 | 4/10 | 4/10 | 3/10 | 2/10 | 3/10 | 2/10 | 3/10 |
|  |  | Diffuse Degeration | --- | --- | --- | --- | --- | --- | --- | --- |

## 4. Discussion

The primary goal of the presented study was to determine the impact of BPA on cardiac specific end points. These analyses are a part of a larger integrated multi-investigator effort performed in parallel with the comprehensive GLP-compliant 2-year CLARITY-BPA chronic exposure study investigating the toxicity of BPA (21, 24). The premise for analysis of these cardiac endpoints was derived from accumulating evidence indicating that the heart is a target for the effects of the endocrine disrupting chemical BPA (16-19, 37). While the heart of both males and females express estrogen receptors (38), the impacts of ER activation by estradiol and the disruptive actions of BPA in the heart are sex specifically regulated and often differ in males and females (12, 13, 17-19). At the onset of the study phenotypic changes related to cardiac remodeling and changes in the collagen extracellular matrix were considered most likely based on the established phenotypes observed in previous mouse studies investigating the impacts of chronic BPA exposures (17-19). In general there was no evidence found here for BPA grossly impacting cardiac endpoints related to hypertrophy in the NCTR-SD rat. Similar to previous findings from the NCTR 90 day BPA toxicity study (25), heart weight was largely unchanged by BPA exposures with increases found only in females exposed to EE. Consistent alterations in LV wall thickness were also not observed. Exposure related changes in collagen accumulation found here were minor and limited to highest EE exposures that resulted in increased collagen accumulation in PND21 males and decreased collagen in hearts of BPA or EE exposed females at PND90 and 6 months. There were higher baseline levels of ventricular collagen in the NCTR-SD rat model compared to the mouse strains used in previous studies (12, 17-19). The relatively higher level of collagen observed in the rat heart is consistent with known species specific differences in the proportions of myocytes and fibroblasts present in murine and rat hearts (39). Whereas the majority of cells in the mouse heart are cardiomyocytes, >60% of the cells in the rat heart are fibroblasts. This difference is likely related to size differences and differences in cardiac physiology and contractile function of mice and rats. For example, the heart rate in mouse is around 700 beats/min, whereas the rate in rat is between 300-400 beats per min. The significant species differences in numerous parameters of cardiac function, structure, cellular makeup and physiological response to pathology require caution when extrapolating observations across different animal models and to human disease.

### 4.1 Exposure to BPA increases cardiac pathology: progressive cardiomyopathy

Rodent PCM is a common background lesion of unknown etiology that is suspected to arise from a localized microvascular dysfunction. The resulting lesions phenotypically progress from minor to extensive focal mononuclear cell infiltration, myocyte degeneration, and fibrosis. The common occurrence of PCM lesions in some rat strains has presented challenges for analyzing cardiotoxicity in regulatory toxicology studies of chemicals and pharmaceuticals due to the difficulty of distinguishing background cardiomyopathy from exposure related effects of chemical exposures (33, 34, 40-43). The Sprague-Dawley rat has been characterized as having an especially high incidence of PCM compared to other rat strains and mice (43).

Due in part to the standardized design of most short and long-term toxicological studies, PCM has not been well evaluated in young animals. Although anecdotal evidence suggests that PCM has been observed in very young rats (44), to our knowledge the analysis presented here is the first quantitative characterization of cardiomyopathy in rats at PND21. A remarkable abundance of early PCM lesions were present in the hearts of most of the young prepubertal study animals. Consistent with PCM found in adults, the lesion incidence and severity was greater in control males than in females, and accordant with BPA have low dose cardiotoxic effects, even at this early age one female in each of the two lowest BPA exposure groups (2.5 and 25 μg/kg/day) and a male in the 250 μg/kg/day group presented with a diffuse degeneration phenotype in which evident pathology involved much of the myocardium was detected (40).

The significance of observing this pathology in these adolescent rats requires additional study to define the nature and etiology because of the relatively limited number of study animals analyzed in each group and the infrequency of this most severe phenotype. The diffuse degeneration phenotype has been taken as indicative of cardiotoxicity (40), thus the findings of this phenotype as early PND21 may suggest that the NCTR-SD strain is partially sensitive to vascular perturbations of the EDC activities of BPA that result in this rare cardiovascular pathology. An increased incidence of the diffuse degeneration myocardial pathology was also observed at PND90 in both BPA and EE exposed male and female from the developmentally exposed and the continuously exposed arms of the study. While the dose response characteristic of this relatively rare phenotype should also be interpreted with caution, it is notable that diffuse degeneration was observed most frequently in the lowest BPA exposure groups (2.5 - 250 μg/kg/day) of the continuously dosed animals.

While complicated by the high level of background pathology, the increases in PCM observed in females at PND21 and the notable increase in myocardial degeneration at PND90 suggests that BPA and EE may impact cardiovascular functions resulting in an early onset of vascular dysfunction and progression of cardiomyopathy. At the 6 month time point there is notable absence of the diffuse degeneration phenotype. It is considered possible that the absence of this phenotype at 6 months was the result of hypertrophic increases in heart size that precluded lesion involvement reaching the grading criteria threshold of >81% LV area involvement to be scored as diffuse degeneration (34-35). The histopathology findings at the 6 month time point (and to a lesser degree at PND21), as well as those for other endpoints, however, should be interpreted cautiously as results may be confounded by the potential for low level BPA contamination in many of the 6 month old animals including nearly all of the control animals analyzed (24). It is notable that the evidence for a low level of background BPA exposure is only inferred from detection of BPA-glucuronide in control and 2.5 μg/kg/day BPA groups from the NCTR BPA 90 day subchronic study (25). The low levels of background BPA detected there were linked to housing of study animals with animals exposed to high concentrations of BPA (24). Because some study animals analyzed here were similarly housed for varying durations in animal rooms with animals dosed at 250,000 μg/kg/day BPA, it is possible that an unintended exposure to low levels of BPA may have occurred. If a similar unintended exposure did occur in the current study, BPA levels in some control animals may be indistinguishable from the 2.5 μg/kg/day BPA exposure groups (26).

### 4.2 Effects of Exposure on Body Weight

Whereas studies using a variety of different developmental exposure paradigms in differing rat and mouse strains have reported obesogenic and diabetogenic actions associated with BPA exposure (reviewed in (45), there were no detectable impacts on body weight found in the study cohort analyzed at any age or dose of BPA or EE. This finding is consistent with numerous previous studies including our own that have consistently found evidence for sex specific changes in cardiovascular and metabolic phenotypes of both rats and mice, but do not detect evidence for BPA causing increases in adiposity and body weight (18, 19, 46-52). These findings are also consistent with the majority of systematic reviews analyzing the strength of evidence from human epidemiologic data which fail to support a causal link between BPA and obesity or type two diabetes (53-55), although a recent systematic review and meta-analysis has found evidence for a link between urinary BPA levels and risk of diabetes and increased obesity (4). The inconsistent findings reported in both experimental animal studies and analyses of human data evaluating the obesogenic potential of BPA is considered most likely due to differences in experimental procedure, experimental design and analysis methods.

The observed significant differences in body weights between continuously dosed males and males dosed only until weaning at PND21 indicate that there were sex specific impacts related to post-weaning dosing procedures and/or the CMC vehicle. The sex-specific decreased weight of males dosed daily with vehicle by gavage is consistent with previous studies showing that prolonged postnatal stress in males decreases weight gain over time, and that female SD rats are resistant to these effects of stress (56-58). Previous studies investigating impacts of prenatal BPA exposure in the developing brain of the NCTR-SD rat have also found that compared to offspring of untreated controls, gavage of pregnant dams with 0.3% CMC resulted in altered estrogen receptor expression in the amygdala of their neonatal offspring (59). Those findings indicated that some endpoints analyzed can be sensitive to either the vehicle and/or maternal stress resulting from the dosing procedures. However, it is not possible to differentiate whether these observed effects were related to different durations of exposure to CMC vehicle, continuous daily restraint and gavage, or their combined effects. Recently chronic oral exposure to CMC in drinking water of mice has been shown to disrupt normal metabolism by altering mucus-microbial interactions that modify gut bacteria composition and metabolic function, effects that caused increased intestinal inflammation, obesity and metabolic syndrome (60). Based on that study, it would be expected that chronic exposure to vehicle would cause an increase in weight. However, it is not possible to rule out the possibility that both exposure to CMC and continuous daily restraint and gavage may be interacting and contributing to the body weight differences observed in males. Determining whether manipulations related to daily gavage or the test material vehicle is responsible for the observed confounding impacts on body weight between continuously or developmentally dosed cohorts will require specific experimental study to define the source(s) of differences in weight gain and possibly other phenotypic impacts.

### 4.3 Study Limitations

Along with confounds related to the duration of dosing, there were additional constraints related to conforming with the CLARITY-BPA consortium study design that did not allow experimental manipulation or direct assessment of changes in cardiac function or cardiovascular endpoints of interest (e.g. contractility and blood pressure) that may have limited sensitivity to detect exposure related phenotypes. Overall previous experimental studies in CD-1 and C57Bl6/n mice found compelling evidence for BPA to alter the collagen extracellular matrix of the heart and to potentially influence cardiac function and negatively impact heart health. The pathology associated with the majority of these effects became most evident following adverse cardiovascular events such as cardiac ischemia or myocardial infarction (17, 19). It is well accepted that experimental interventions resulting in increased β-adrenergic stress, ischemic injury or genetic manipulations are often necessary to reveal cardiac fibrosis and hypertrophy or phenotypes indicative of overt cardiac pathology in rodent models (61). Such manipulations were not possible in this study and only post mortem tissues were available for analysis. The inability to experimentally manipulate study animals is considered a limitation as the morphometric endpoints we were able to analyze are relatively insensitive phenotypes. Additional studies using procedures or models that develop CV disease phenotypes in rat could be useful for clarification of BPA impacts on the heart.

## 5. Conclusions

Largest observed morphometric effects were due to treatment duration which altered body weight and cardiac collagen accumulation. Overall, neither BPA nor EE caused hypertrophy or overtly altered fibrosis in the NCTR-SD rat at PND21, PND90 or PND180. However, compared to CD-1 and C57Bl6/n mice the NCTR-SD rat has higher baseline collagen levels and a high level of degenerative cardiomyopathy which may have limited the ability to detect exposure-related impacts on these end-points. Exposures to either BPA or EE increased incidence and severity of progressive cardiomyopathy in females at PND21, and increased the severity of cardiomyopathy in both sexes at PND90. Increases in PCM are indicative of modest exposure related cardiotoxicity that may be the result of an increase in adverse vascular events.

## Acknowledgments

We are thankful for the outstanding work of Drs K. Barry Delclos, Luisa Camacho, and the staff of the National Center for Toxicological Research/Food and Drug Administration, Thaddeus Schug and Retha Newbold of NIEHS on the CLARITY-BPA and Dr. Jennifer Fostel, Cari Martini and the Chemical Effects in Biological Systems (CEBS) team from the NIEHS/National Toxicology Program. The encouragement and valuable advice of Drs John Meitzen and Heather Patisaul during the writing of this manuscript was instrumental in its completion.

